# Integrative biology defines novel biomarkers of resistance to strongylid infection in horses

**DOI:** 10.1101/2021.04.26.441388

**Authors:** Guillaume Sallé, Cécile Canlet, Jacques Cortet, Christine Koch, Joshua Malsa, Fabrice Reigner, Mickaël Riou, Noémie Perrot, Alexandra Blanchard, Nuria Mach

## Abstract

The widespread failure of anthelmintic drugs against nematodes of veterinary interest requires novel control strategies. Selective treatment of the most susceptible individuals could reduce drug selection pressure but requires appropriate biomarkers of the intrinsic susceptibility potential. To date, this has been missing in livestock species. Here, we selected Welsh ponies with divergent intrinsic susceptibility to cyathostomin infection and found that their potential was sustained across a 10-year time window. Using this unique set of individuals, we monitored variations in their blood cell populations, plasma metabolites and faecal microbiota over a grazing season to isolate core differences between their respective responses under worm-free or natural infection conditions. Our analyses identified the concomitant rise in plasmatic phenylalanine level and faecal *Prevotella* abundance and the reduction in circulating monocyte counts as biomarkers of the need for drug treatment. This biological signal was replicated in other independent populations. We also unravelled an immunometabolic network encompassing plasmatic beta-hydroxybutyrate level, short-chain fatty acid producing bacteria and circulating neutrophils that forms the discriminant baseline between susceptible and resistant individuals. Altogether our observations open new perspectives on the susceptibility of equids to cyathostomin infection and leave scope for both new biomarkers of infection and nutritional intervention.

## Introduction

Infection by gastro-intestinal nematodes is a major burden for human development worldwide as they both affect human health^1^ and impede on livestock production^2^. Worldwide reports of anthelmintic drug failures against nematodes of veterinary importance have accumulated^3^, threatening the sustainability of livestock farming in some areas. The same pattern applies in horses whereby widespread benzimidazole failure and intermediate pyrantel efficacy against cyathostomin populations have been reported^4–6^. These small strongyles locate in their host hindgut and are responsible for growth retardation in young animals^7,8^. The massive emergence of developing larval stages from the caeco-colic mucosa can cause a larval cyathostominosis syndrome^9^ that remains a leading cause of parasite-mediated death^10^.

Factors contributing most to the selection of drug-resistant cyathostomin populations in equids remain uncertain^4,5^. However, significant and heritable inter-individual variation in resistance to strongylid infection has been reported in both domestic^11,12^ and wild horse populations^13^. This variation leaves scope for restricting drug application to the most susceptible horses, thereby alleviating the selection pressure on parasite populations. To date, the genetic architecture of this trait has not been defined in equids, although indications from ruminant species would be in favour of a polygenic architecture^14–16^ defining a stronger type 2 cytokinic polarization in resistant individuals^17,18^.

Identifying biomarkers of this intrinsic resistance potential would both contribute to understanding the host-parasite relationship and to defining relevant biomarkers for use in the field. Current targeted-selective treatment schemes are based on faecal egg count (FEC) that has suboptimal sensitivity and remains time-consuming despite recent advances that should ease egg detection^19^. As a result, its uptake in the field varies widely across countries and remains limited^5,20^ despite being cost-effective^21,22^. To date, limited alternative biomarkers have been identified. Alteration in serum albumin level and decrease in circulating fructosamine were the main features found in cyathostomin infected ponies^23^. Independent observations concluded that mixed strongyle infection was associated with mild inflammatory perturbations^24^. We previously highlighted that susceptible ponies had lower monocyte but higher lymphocyte counts than resistant individuals upon natural strongylid infection^25^. More susceptible individuals also exhibited differential modulation of their faecal microbiota, including enrichment for the *Ruminococcus* genera^25^, corroborating independent observations of alterations in the gut microbiota composition of infected horses^25–28^.

In horses as in other host-parasite systems, limited efforts have been made to isolate compositional shifts in plasma metabolites following parasite nematode infection. Beyond murine models of helminth infection^29,30^, implementation of this technology could define a urinary biomarker of infection by *Onchocerca volvulus* in humans^31^. This was however not reproduced in other cohorts of patients^32^. In livestock species, a single study has applied metabolomic profiling on horse faecal matter to identify biomarkers of infection by parasitic nematodes but found little differences between horses with contrasted levels of strongylid infection^26^.

In any case, these observations remain limited to individual host compartments and do not provide an integrated perspective of the physiological underpinnings associated with susceptibility to infection. Because we are aiming to distinguish between individuals before the onset of degraded clinical signs, the biological signals may be subtle. To this respect, integration of systems biology data - that consider multiple high-dimensional measures from various host compartments - is expected to better identify the multiple features defining a given physiological state^33^.

Under this assumption, we combined and analysed metabolomic, metagenomic and clinical data collected on a selected set of intrinsically resistant and susceptible ponies to identify the physiological components underpinning their resistance potential. Our data defined a strongylid infection signature built around lower circulating monocytes, enriched plasmatic phenylalanine concentration and higher *Prevotella* load in faecal microbiota that we could replicate in independent populations. We also identified an immunometabolic signature centered on neutrophils that best discriminated between resistant and susceptible individuals across strongyle-free or natural infection conditions. These results begin to define the physiological bases supporting the intrinsic resistance potential to strongylid infection in equids.

## Results

This experiment was based on a set of individual Welsh ponies with divergent resistance potential to strongylid infection. During the experiment (2015), we produced metabolomic data that we present herein and integrated this plasma-related dataset with previously described faecal bacteria profiles and clinical parameters^25^. We analysed the data produced for each group under worm-free conditions (day 0) or following natural infection (day 132) to identify biomarkers of infection, ii) establish a holistic view of the physiological underpinnings of the resistance potential to strongyle infection in horses.

### 1. Pony divergence toward strongyle infection is significant and sustained

We selected 20 female Welsh ponies with divergent susceptibility to strongylid infection. Their susceptibility potential was predicted from their past FEC history (at least three FEC records over two years between 2010 and 2015). During the experiment (2015) and following natural infection (day 132), 17 ponies displayed FEC values in good agreement with their predicted potential. In that case, 8 of the susceptible ponies (**TS**) had FEC above the considered 200 eggs/g cut-off for treatment (average FEC = 419 ± 149 eggs/g) at day 132, and 9 resistant ponies (**TR**) were below this threshold (average FEC = 56 ± 77 eggs/g). Our prediction hence achieved an accuracy of 85%. Other ponies that did not match expectations had either higher susceptibility (850 eggs/g for the predicted resistant individual) or too low FEC (0 and 50 eggs/g for the two susceptible individuals).

In agreement with this observation, FEC measured in the TS and TR ponies during the five years preceding the experiment departed significantly from the herd mean (0.6 standard deviations; *P* = 0.01 and *P* = 0.02 for TS and TR groups respectively; Fig 1). To ensure that their intrinsic potential was true, we compared their FEC records collected after the experiment took place (between 2015 and 2020) to that of their herd (1,436 individual records). We confirmed that their divergence was sustained (−0.76 and +0.63 standard deviation from mean in TS and TR ponies respectively; *P* = 0.02 in both cases) throughout the following years (Fig 1), hence validating their intrinsic potential. This corresponded to an average FEC of 43 eggs/g (min = 0; max = 2,200 eggs/g) and 756 eggs/g (min = 0; max = 2,800 eggs/g) in TR and TS ponies respectively, while the average herd FEC was 320 eggs/g (min = 0; max = 5,400 eggs/g).

**Figure 1.**
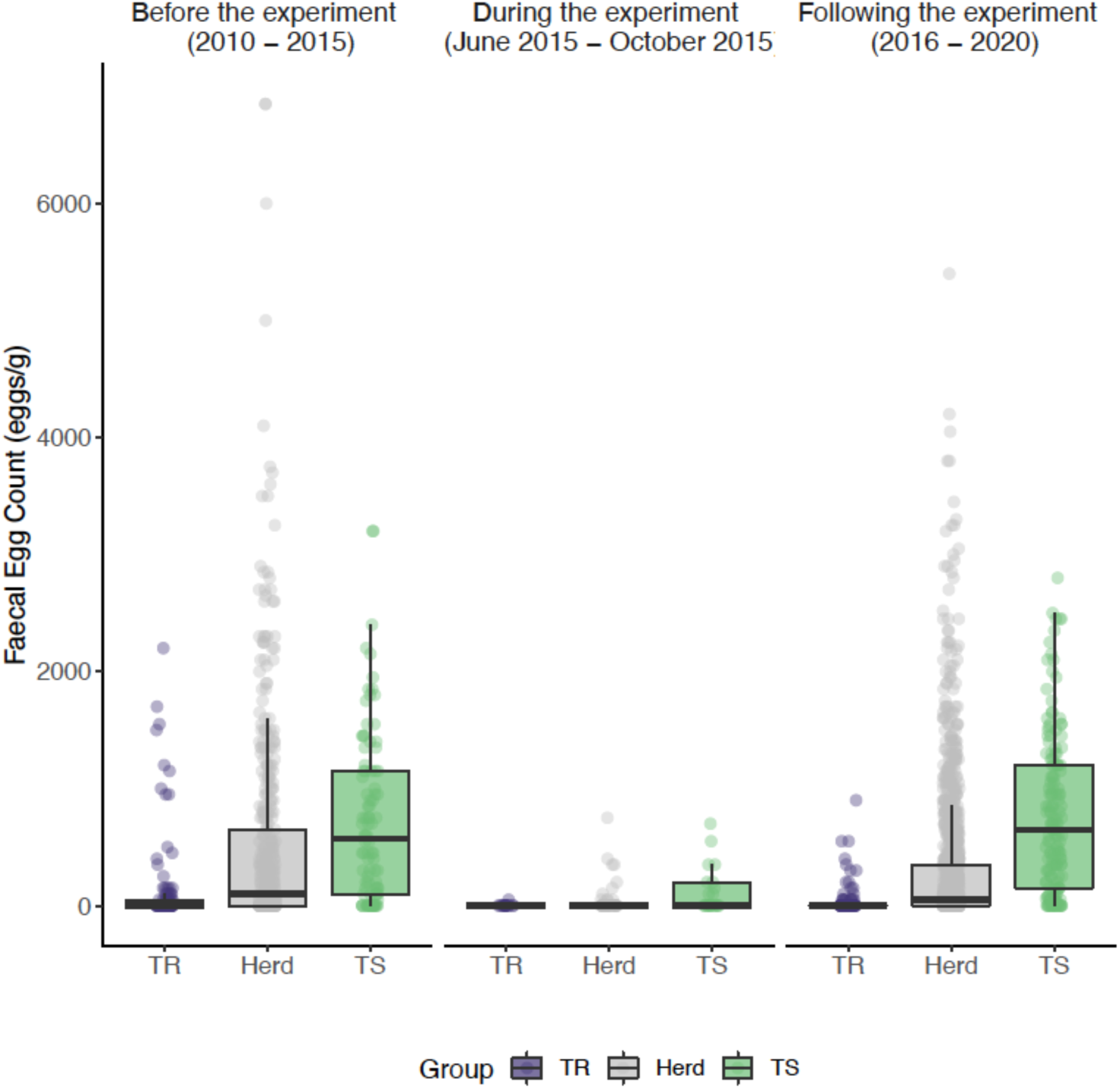
FEC-informed pony potential prediction is robust through time. The distribution of observed Faecal Egg Counts before the experiment (2010-2015), during the experiment (2015) and after the experiment (2015-2020) in the predicted susceptible (8 individuals, 271 records; green) and resistant (9 individuals, 223 records; purple) ponies and their herd unselected counterparts (127 individuals, 1,436 measures; grey) is represented. Dots stand for individual measures and boxplots represent the data distribution (mean materialized by a vertical bar within the box that stands for the 25% to 75% interquartile range).

### 2. Metabolomic profiling highlights the association between plasmatic phenylalanine level and FEC

To identify markers of pony intrinsic resistance potential, we measured variation in their plasma metabolites using ^1^H-NMR between worm-free conditions (day 0) and after natural infection (day 132). This metabolomic profiling of TR and TS ponies throughout the grazing seasons found a total of 791 metabolic buckets, corresponding to 119 unique metabolite signals (Fig 2, supplementary Table 1). These included several amino acids, energy metabolism-related metabolites, saccharides, unknown lipids, and organic osmolytes in the plasma. Among these signals, we observed 29 unassigned bins in three main windows ranging between 1.115 and 1.435 ppm, 3.385 and 4.305 ppm or 6.805 and 7.895 ppm (supplementary Table 1).

**Figure 2.**
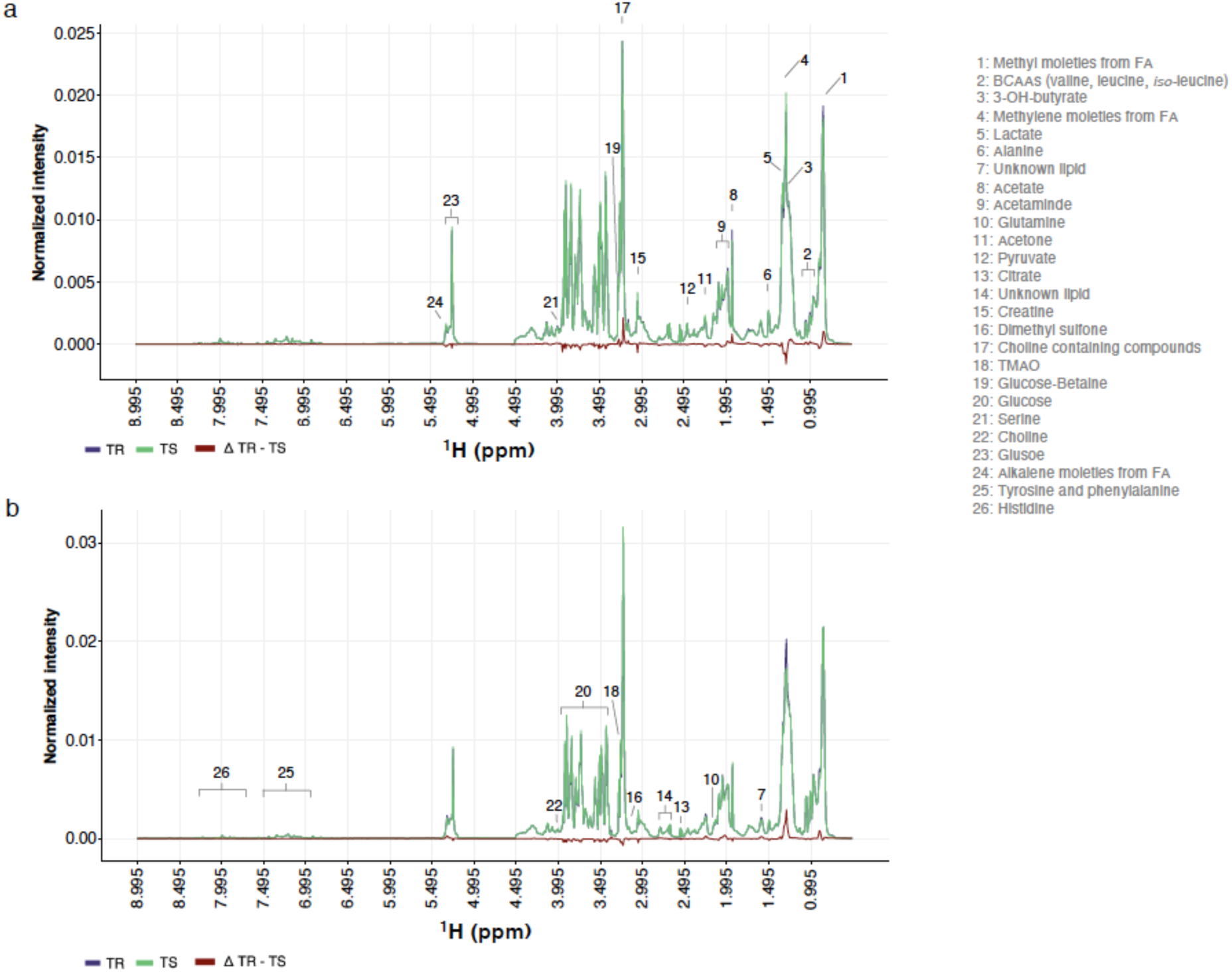
Representative ^1^H-NMR spectra measured in resistant and susceptible ponies. Group average ^1^H-NMR signal intensities are plotted against the considered chemical shifts ranging from 0.995 to 8.995 ppm and overlaid for resistant (TR; purple) and susceptible (TS; green) pony groups at day 0 (strongyle-free; panel a) or day 132 (strongyle infected; panel b). Differential intensity between groups is drawn in red. Associated metabolites are annotated by numbers.

The ANOVA – simultaneous component analysis (ASCA) applied to metabolomic time series data identified significant temporal variation (*P* < 0.001) but no differential rewiring occurred between TR and TS pony metabolomes (*P* = 0.54). The temporal variation was structured around five signals (supplementary Fig 1) associated with alkalene moieties from lipids (^1^H-NMR signal at 5.265−5.355 ppm), sugar moieties of α- and β-glucose (3.455−3.555 ppm) in overlap with proline (3.395−3.445 ppm) and branched-chain amino-acids (BCAAs) such as valine (1.045−1.055 ppm) and leucine (0.965-0.975 ppm). These signals showed mean leverage of 3.4% (ranging between 1.5% and 7.8%) and squared prediction error below 2.2 x 10^-5^. Among these, the high-intensity signals ascribable to glucose decreased between the strongyle-free (day 0) and strongyle-infected (day 132) conditions (supplementary Fig 1).

Despite the lack of systematic modifications of metabolomes between TR and TS ponies, we sought to identify individual metabolites that would reflect the intrinsic susceptibility status of ponies before any infection took place (day 0) or metabolites that would differentiate individuals in need of treatment at the end of the grazing season (day 132). Considering the nominal *P*-values of 5%, we identified differences in dimethyl-sulfone and lysine associated signals that were all decreased in the TS pony group (supplementary Fig 2a). None of these signals were however significantly correlated with final FEC measured at day 132 (Spearman’s ρ ranging between -0.08 and 0.34, supplementary Fig 2b).

The same approach applied to TR and TS ponies after natural infection at day 132 found lower levels of phenylalanine (^1^H-NMR signals at 3.285-3.305 ppm and 7.345-7.445 ppm) in TR compared to TS ponies (nominal P-value = 6 x 10^-3^). In addition, an unidentified metabolite between 6.815-6.815 ppm (U13) was significantly lower in TR (*P* = 0.04). Phenylalanine signal intensities increased with FEC (Spearman’s ρ ranging between 0.55 and 0.60, *P*< 0.05, n = 17). Using the faecal metabolomic data from another independent set of British horses ^26^, we could validate this signal. In that study and in line with our results, faecal phenylalanine level was significantly increased in horses with higher FEC (*Wilcoxon’s test = 27*, *P-value* = 0.02, Fig 3C).

**Figure 3.**
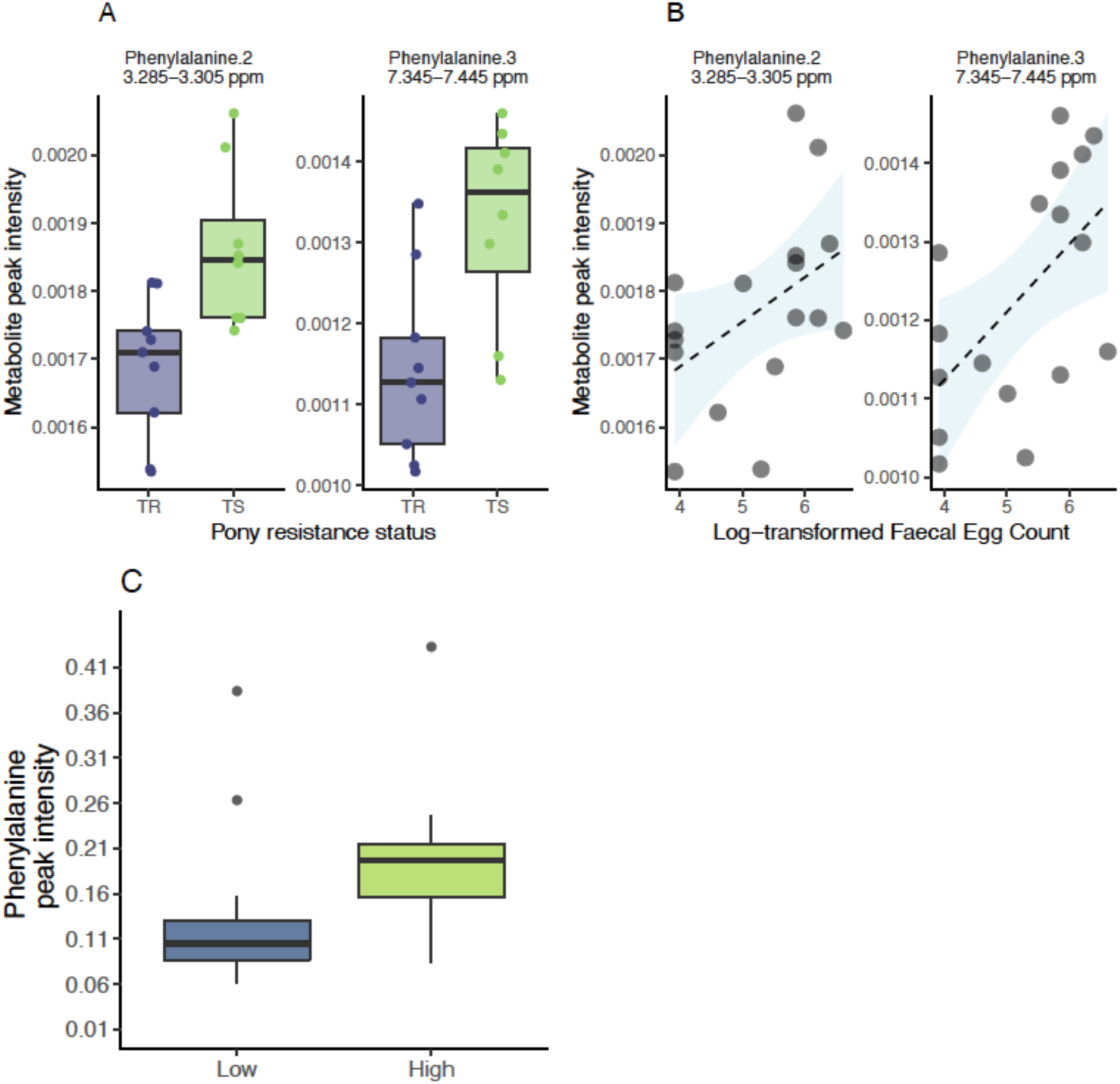
Differential metabolites between infected resistant and susceptible ponies (day 132) Panel A shows metabolite signal intensity distribution in each pony susceptibility group (purple: resistant, TR; green: susceptible, TS) at day 132. Panel B shows the relationship between these metabolite signal intensities (X-axis) at day 0, and matching log-transformed Faecal Egg Count (Y-axis) at day 132. Panel C describes observed phenylalanine levels in the faecal matter of an independent cohort of British horses with low or high FEC ^26^.

Altogether these results indicate that strongylid infection experienced by TR and TS ponies did not induce metabolome-wide modifications. However, phenylalanine was the most discriminant between TR and TS ponies under infection and stands as a biomarker of the need for treatment.

### 3. Multi-compartment data integration identifies circulating monocytes, phenylalanine and faecal *Prevotella* levels as discriminant features under infection

To mine the physiological differences between infected TR and TS ponies deeper, we applied a data integration framework (sGCC-DA) bringing together clinical, metabolomic and previously analyzed faecal microbiota data from these individuals ^25^. The first component of the sGCC-DA better discriminated between TR and TS ponies and retained nine clinical parameters, 15 bacterial genera and ten plasma metabolite signals (Fig 3, supplementary Figs 3 and 4).

The network of correlations between these features was structured around two major clusters (Fig 4, supplementary Figs 3 and 4). A first core of features built around average daily gain (ADG) and four commensal gut bacterial genera (*Anaerovibrio, Fibrobacter, Prevotella,* and *Treponema*) displayed higher levels in TR ponies under infection (Fig 4, supplementary Figs 3 and 4). These features were negatively correlated to a strongylid susceptibility-associated cluster that encompassed neutrophil counts (2.8 ± 0.51 and 3.05 ± 0.8 million cells/mm^3^ in TR and TS ponies), the haematocrit (average of 38.74% ± 1.7 vs. 41.45% ± 4.1 in TR and TS ponies) and the plasmatic levels of serine and essential amino acids such as phenylalanine, lysine and tryptophan (Fig 4, supplementary Figs 3 and 4). This underscores the association between phenylalanine and FEC (Fig 3b, c).

**Figure 4.**
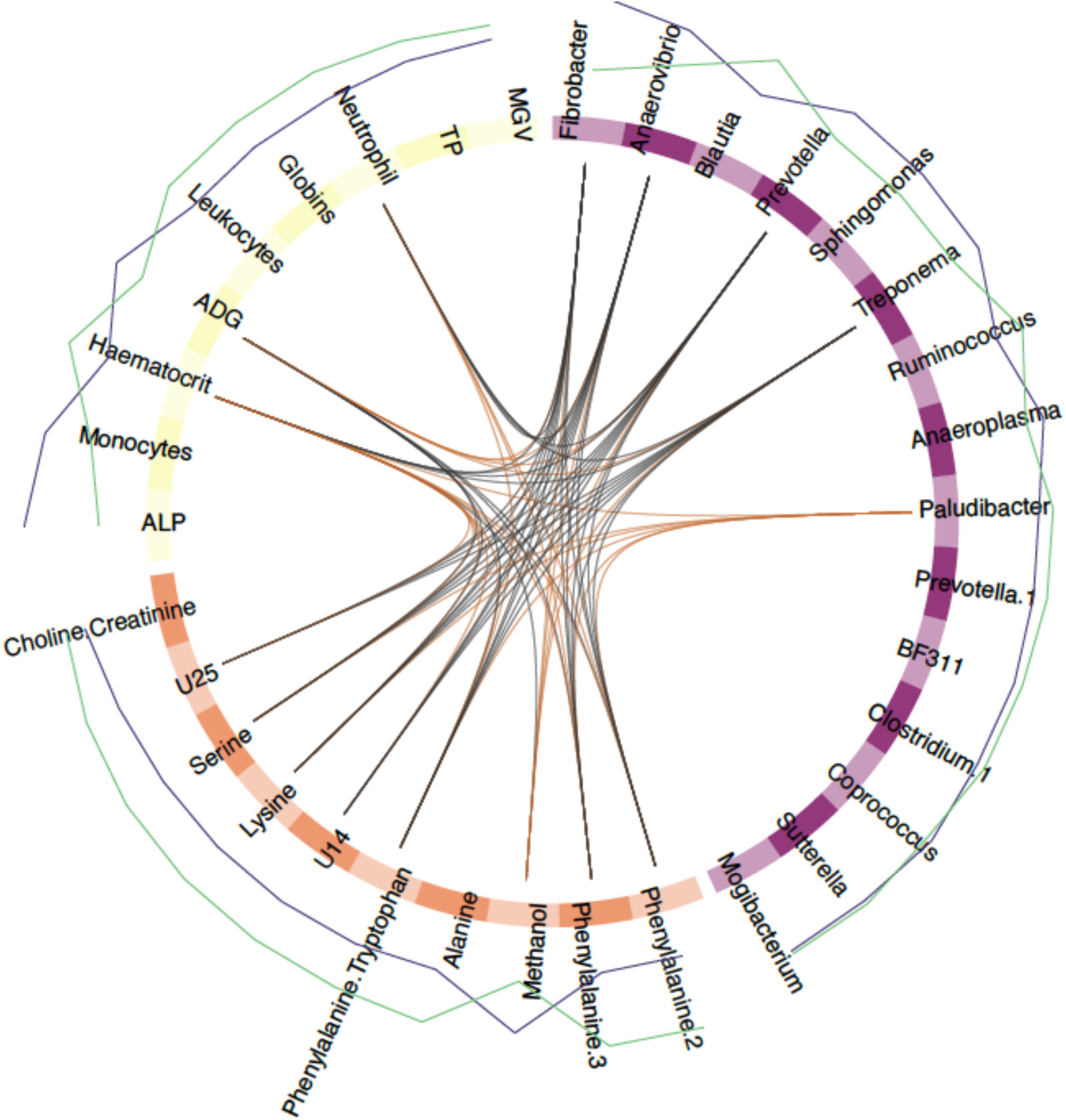
Circos plot showing features best discriminating between infected resistant and susceptible ponies (day 132) For each of the three input data types (clinical data in yellow, bacterial genera in purple and metabolite signals in orange), the features best discriminating between resistant and susceptible ponies are listed. A link is materialized between two features if their shared correlation is above 0.45 (chocolate if positive, grey otherwise). External green and purple lines represent the relative feature level in each pony susceptibility group (resistant, TR: purple; susceptible, TS: green). MGV: Mean Globular Volume; TP: Total Protein; ALP: Alkaline Phosphatase; U14, U25: Unknown metabolites 14 and 25.

Additional features included higher monocyte counts (1.178 ± 0.23 and 1.038 ± 0.24 million cells/mm^3^ in TR and TS ponies on average) and higher levels of plasmatic alkaline phosphatase (ALP; 5.39 ± 0.15 and 5.25 ± 0.12 Units/L) in TR ponies (Fig 4). But these parameters displayed least covariation with other parameters (Fig 4).

Analysis for KEGG pathway enrichment of metabolite signals found significant over-representation of Aminoacyl-tRNA biosynthesis (FDR = 1.1 x 10^-4^) underpinned by the presence of alanine, lysine, phenylalanine, serine, and tryptophan (supplementary Table 2). Of note, alanine and phenylalanine plasmatic levels defined significant enrichment for the dengue fever (supplementary Table 2). The most significant enrichment was defined by the *Blautia*, *Coprococcus* and *Ruminococcus* association found in forms of pediatric Crohn’s disease (FDR = 2.5 x 10^-3^). Two other significant enrichments included infection-mediated perturbations associated with the HIV-1 virus in humans (underpinned by *Anaerovibrio* and *Clostridium* genera, FDR = 1.12 x 10^-4^) or with murine model of *Plasmodium* infection (underpinned by *Anaeroplasma* and *Clostridium*, FDR = 0.05).

To validate the association of the most discriminant blood and bacterial features with FEC, we used either a previously published horse data set ^26^ for bacterial count, or additional blood samples taken from 25 strongylid infected ponies in 2020 (Fig 5). Out of the five most discriminating bacterial genera identified by our sGCC-DA approach under infection at day 132, *Prevotella* (FDR = 0.19; nominal *P-value* = 0.04) also showed significant differences in their mean abundances between Peachey et al.’s horses with low or higher FEC (Fig 5). Monocyte counts were also negatively correlated with FEC levels in the independent set of ponies (Spearman’s ρ = -0.31, *P-value* = 0.14; Fig 5).

**Figure 5.**
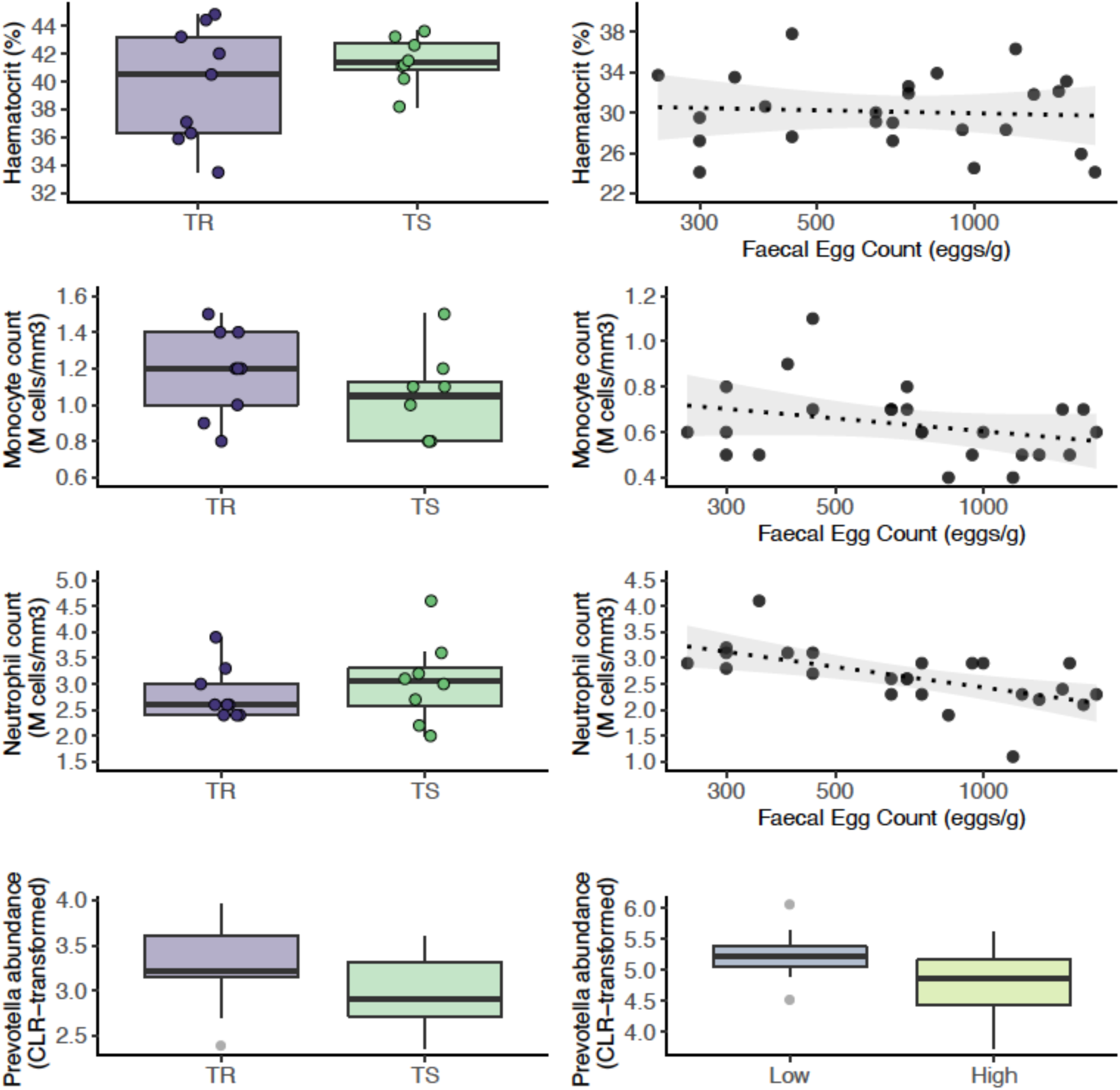
Validation of the most discriminant features between resistant and susceptible ponies under strongylid infection. Each row presents the feature level in resistant and susceptible ponies (left panels) and the association with Faecal Egg Count recorded in an independent set of ponies or horses (for the *Prevotella* genus). CLR: Centered Log-Transformed. The figure highlights the significant positive relationship between circulating monocyte count and faecal abundance of the *Prevotella* genus in independent individuals.

The trend was however opposite for the neutrophil population (Spearman’s ρ = -0.63, *P-value* = 9 x 10^-4^; Fig 5) and no relationship between FEC and haematocrit was found in this independ set of individuals (Spearman’s ρ = -0.07, *P-value* = 0.7; Fig 5).

These findings hence retain increased *Prevotella* abundance in faecal material as a conserved signal in equids with reduced FEC, and monocyte counts appears to be a good predictor of FEC level.

### 4. Short-chain fatty acid producing bacteria, plasmatic lysine and circulating neutrophils recapitulate pony intrinsic potential across conditions

Using the same analytical framework, we aimed to identify features that would best define the intrinsic pony resistance potential across worm-free or natural infection conditions. The sGCC-DA applied on records measured under worm-free conditions (day 0) identified reduced circulating neutrophil and leukocyte counts in the TR ponies as the most discriminant clinical parameters (supplementary Figs 5, 6, and 7). Cell counts of these two populations were tightly linked with plasmatic levels of 1-methylhistidine (7.785 ppm; reduced in TR ponies), lysine (1.445-1.465 and 1.845-1.915 ppm; increased in TR ponies) and β-hydroxybutyrate (4.125-4.135 ppm; increased in TR ponies). In addition to these parameters, maximal covariance was obtained for a few genera from the Actinomycetia class (order Corynebacteriales), namely *Dietzia*, *Gordonia*, *Mycobacterium* and *Sacharopolyspora* (order Pseudonocardiales), that all showed higher relative abundances in resistant ponies, as well as the candidate genera BF311 within Bacteroidetes (e.g., *BF311*; supplementary Figs 5). On the opposite, higher relative abundance of butyrate-producing Clostridia, specifically *Anaerofustis, Coprococcus* and *Ruminococcus* were found in TS ponies (supplementary Figs 5). These genera showed positive correlations with circulating lymphocytes and neutrophil counts (ranging between 0.51 and 0.68 for lymphocyte counts and between 0.38 and 0.51 for neutrophil counts respectively; supplementary Figs 5 and 7). Of note, the presence of *Prevotella* and *Desulfovibrio* among the set of covarying features defined significant enrichments compatible with *Plasmodium* infection in mice or *Schistosoma haematobium* infection in humans (FDR = 0.04 in both cases; supplementary Table 2).

Combining these differential features between pony groups under worm-free conditions with that found after strongylid infection retained a core set of seven markers consistently discriminating TR and TS ponies across infection conditions (supplementary Fig 8). For these markers, we aimed to identify differential trends between both groups across infection conditions.

First, circulating neutrophil and leukocyte counts showed significant increase following infection (*P* = 0.02 and 4 x 10^-4^ respectively), but this increase was not different between TR and TS ponies (*P* = 0.5 in both cases; supplementary Fig 8 and supplementary Table 3). This trend was also not corroborated in another independent set of individuals (Fig 5).

Second, plasmatic lysine level was significantly different between infected TR and TS ponies (*P* = 0.002) and matched independent observations made in faecal samples from another cohort of British horses (Fig 6). Our temporal records also supported a differential decrease of plasmatic lysine levels between TR and TS ponies from worm-free to natural infection conditions (*P* = 0.02).

**Figure 6.**
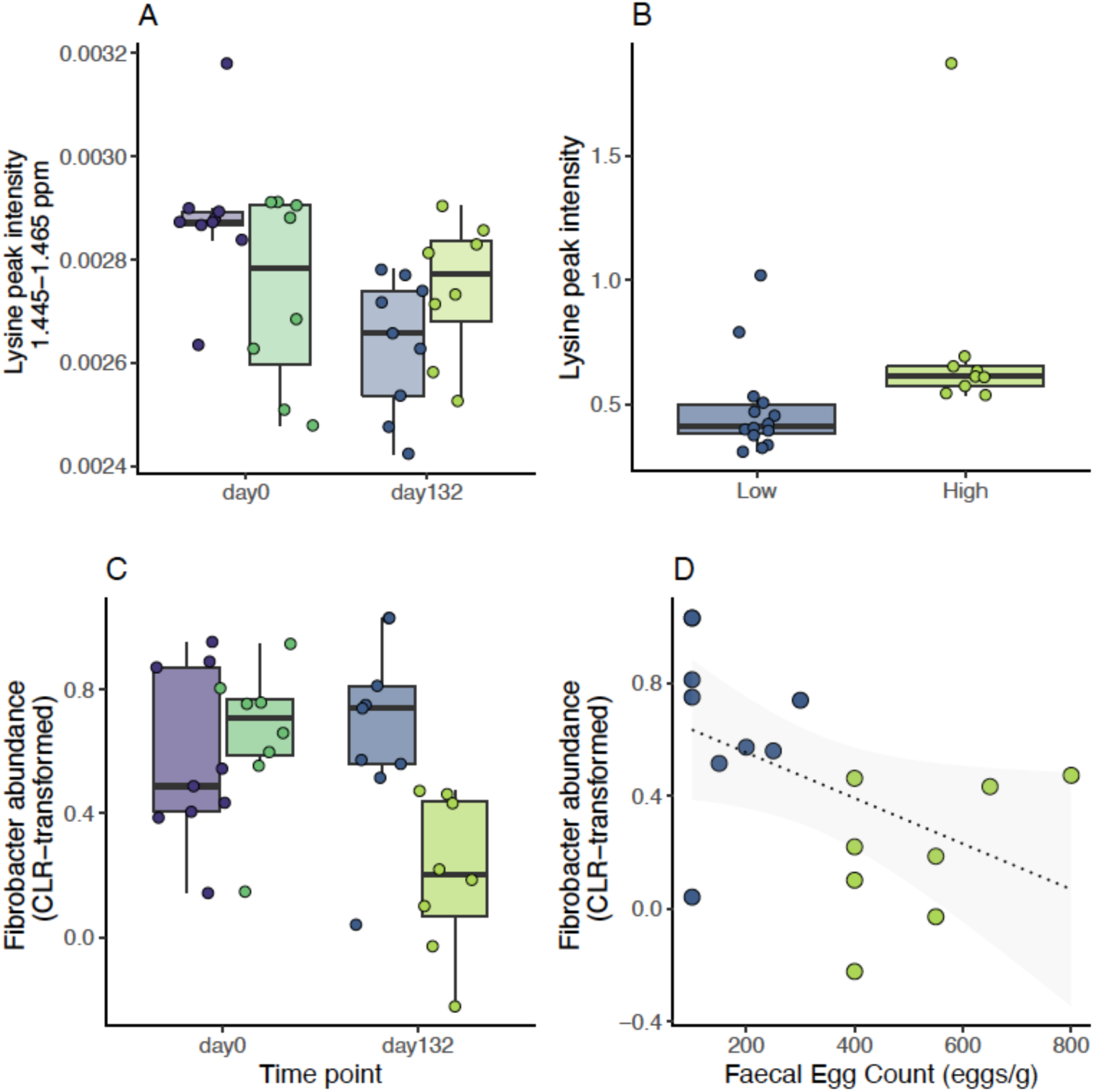
Discriminant features between resistant and susceptible ponies across infection conditions. Measured values of lysine signal intensity (A) and Fibrobacter abundance (C) are plotted across susceptibility groups (TR: resistant, in blue; TS: susceptible, in green) and time points (day0: before infection; day132: under strongylid infection, 132 days after the onset of grazing season). Lysine intensity measured on faecal material in an independent cohort of British horses with low or high Faecal Egg Count is represented in panel B. Panel D shows the relationship between Fibrobacter count in TR and TS ponies at day 132 and measured FEC.

Third, sGCC-DA recognized three dominant gut bacteria, namely *Fibrobacter, Ruminococcus,* and *Treponema*, and one rare microbial taxa (*Coprococcus*) as keystone species whose covariation formed a distinct signature between the two pony groups (Fig 6, supplementary Fig 8). Using linear regression modeling, we found a significant decrease in faecal abundance of *Treponema* (*P* = 1.7 x 10^-2^) upon infection that was shared across pony groups (supplementary Fig 8). On the contrary, the observed rise in *Fibrobacter* abundance at day 132 was slightly milder in the TS ponies (−0.73 ± 0.26, FDR = 4.4 x 10^-2^; Fig 4, supplementary Table 3).

Lysine and *Fibrobacter* would hence mark the intrinsically higher sensitivity of TS ponies.

## Discussion

Inter-individual variation in susceptibility to strongylid infection offers the opportunity to reduce drug selection pressure by selectively treating the most susceptible individuals. To date, the physiological markers associated with this resistance potential are to be defined. Here, we selected resistant and susceptible ponies and confirmed that their potential was sustained over a 10-year time window. Using this unique set of individuals, we performed the first detailed investigations of the variations occurring in three different compartments (blood, plasma and faeces) to identify the underpinnings of their differential resistance to strongylid. Our findings are two-fold. First, we found support for defining phenylalanine, circulating monocytes and faecal abundance of *Prevotella* as biomarkers of strongylid infection in equids. Second, we identified an immunometabolic network encompassing beta-hydroxybutyrate level, short-chain fatty acid producing bacteria and circulating neutrophils that recapitulates the intrinsic resistance potential of ponies under both worm-free and natural infection conditions.

Elevation of plasmatic phenylalanine concentration has long been recognized as a metabolic consequence of bacterial and viral infections^34^. Similar observation has also been made in the faecal matter of infected horses^26^ and we could replicate it from plasmatic samples. Of note, our data highlighted the covariation of this essential amino acid with average daily gain and the concomitant increase in plasmatic alkaline phosphatase concentration. This is compatible with strongyle infection reducing muscle protein synthesis (in line with observed reduced average daily gain), thereby increasing the extracellular release of phenylalanine and its subsequent uptake by the liver (whose activity was tracked by the increased alkaline phosphatase level). This model would match the theoretical framework derived from other infectious processes^34^. Altogether, plasmatic phenylalanine represents another diagnostic option for strongylid infection. The sensitivity and specificity of this marker remains to be determined in a larger cohort for field use in association with monocyte count and faecal *Prevotella* abundance.

The concomitant increase in the plasmatic level of this aromatic amino acid with the monocyte count decrease upon infection in susceptible individuals also opens new perspectives on the pathophysiology associated with strongylid infection in equids. Indeed, observations in humans demonstrated that D-phenyllactic acid - an antibacterial compound derived from phenylalanine and produced by the gut microflora - can activate the hominid-specific Hydroxy-Carboxylic Acid (HCA) 3 G-protein coupled receptor expressed by monocytes, thereby favoring their recruitment^35^. While the only lactate-associated HCA1 receptor exists in equids, the possibility remains that phenylalanine derivatives could recruit immune cells^36^. Under this speculative model, serum phenylalanine (increased in infected individuals), would enter the gut lumen before subsequent transformation into phenyllactic acid by gut bacteria. Bacteria-derived phenyllactic would then be absorbed into the portal vein and serve as a recruitment signal for circulating monocytes. Further studies on this receptor would confirm its role in any differential immunometabolic regulations between the resistant and susceptible pony groups.

Under infection, susceptible individuals exhibited decreased abundance of faecal *Prevotella*. Effects of the *Prevotella* genus are still debated^37^. Experimental colonization of germ-free mice by *Prevotella* promoted the decrease of IL-18 interleukin expression, neutrophil recruitment at the site of infection and gut inflammation^37^. *Prevotella* was also associated with susceptibility to *Plasmodium* infection in mice^38^, increased in patients infected by the human trematode *S. haematobium*^39^ and in the colon of pigs infected by *Trichuris suis*^40^. A *Prevotella*-led inflammatory state would hence define the reduced strongyle infection observed in resistant ponies. This would contradict past findings in mice showing the detrimental association between IL-18 and *Trichuris muris* infection^41^. Because this cytokine seems to be sensitive to its environment, there is scope for microbiota-based regulations^42^ that would differ between resistant and susceptible individuals.

In our attempt to discriminate between resistant and susceptible ponies, we identified a feature network with contrasted proinflammatory abilities. First, resistant ponies exhibited higher plasmatic lysine levels across conditions. Lysine is an essential amino acid for metabolism and immune response. Although the role of its derivative on the immune system is yet to be clarified^43^, it is a natural ligand of the GPRC6 receptor^44^ that can modulate Th-2 response and antibody production by B cells^45^. Second, butyrate-producing bacteria and other bacteria able to metabolize β-hydroxybutyrate into butyrate (e.g. *Coprococcus*) defined differential baselines between pony susceptibility groups under worm-free conditions. The former is known to favour an anti-inflammatory state by inhibiting the neutrophil inflammasome and the release of pro-inflammatory cytokines like IL-1β and IL-18^46,47^. Butyrate binds free fatty acid receptors, like FFAR2 that is enriched on neutrophil cell surface^36^, thereby promoting their intestinal recruitment^48^. Third, faecal *Fibrobacter* was a core discriminating feature of the intrinsic resistance potential. The drastic reduction of faecal *Fibrobacter* abundance in susceptible ponies upon strongylid infection mirrored past observations made in pigs infected with *T. suis,* for which the reduction occurred irrespective of the worm load^49^. In goats, abundance of this genus was negatively correlated with the pro-inflammatory cytokines TNFɑ that is produced by monocytes^50^. *Fibrobacter* bacteria are key cellulose degraders that produce succinate subsequently converted into propionate, another short-chain fatty acid able to modulate the recruitment of monocytes and neutrophils^36^. The reduction of this bacterial genera could hence result in a pro-inflammatory state favourable to the development of strongyle infection in susceptible ponies.

This immuno-metabolic network tying the host metabolites with its gut microbiota also points towards a key role of neutrophils in the definition of strongylid susceptibility. In line with this strand of evidence, we found a consistent covariation between circulating neutrophils and intrinsic susceptibility to strongylid infection. This relationship was however not validated in another independent population with higher excretion levels than that found during our 2015 experiment. The role played by neutrophils hence remains to be fully characterized as for other host-helminth models^51,52^. Transient neutrophilia (in the range of 9 x 10^9^ cells per L) was previously described as the only haematological variation in ponies subjected to experimental infection by cyathostomins^23^. But neutrophils are not typically associated with type 2 immunity - that is effective against helminths^53^ - and they do not play a role against the infection by *Trichinella spiralis*, a clade I nematode^54^. However, neutrophils can cooperate with macrophages to bring nematode infection under control as found for *Litomosoides sigmodontis*^55^, *Strongyloides stercoralis*^56^ and *Heligmosomoides polygyrus*^57^. They also appear to be regulated by the type-2 cytokinic environment to prevent damages to the host^51^. In horses, neutrophils respond to IL-4 stimulation - a type 2 cytokine - by an increase in the pro-inflammatory cytokines TNF-ɑ and IL-8 but a decrease in IL1-β^58^ that could be detrimental to the anti-helminth response. The covariation between neutrophil counts and the enhanced susceptibility in horses may thus result from qualitative differences. Application of single-cell RNAseq on neutrophil populations from the two pony backgrounds could provide significant advances in the understanding of their respective properties. Neutrophils were however not part of the recently released equine mononuclear cells atlas^59^. Quantification of cytokines and reactive oxidative species production after *in vitro* exposure to infective strongylid larvae could also unravel distinct properties and putative bias toward a more effective response in resistant individuals.

Overall, this work suggests that FEC records from at least two years are sufficient to define the resistance potential of an equid to strongyle infection. It also proposes phenylalanine, monocyte counts and faecal *Prevotella* abundance as biomarkers of infection. Last, we identify a neutrophil-centered network tying together gut microbiota members and a few metabolites as the discriminant baseline between susceptible and resistant individuals. These results open novel perspectives for the understanding of strongylid susceptibility in equids and for nutritional modulation of these infections.

## Materials and methods

This study relied on measurements previously gathered during the one described previously^25^. Measured features are summarized herein and additional ^1^H-NMR and FEC parameters collected during this experiment are described. Data were analysed with R version 4.0.2 unless stated otherwise.

### Selection and monitoring of 20 ponies with divergent susceptibility profile to strongyle infection

Ponies were selected from the experimental herd according to their FEC history recorded since 2010. Briefly, individual pony random effect was estimated from individual records after correction for environmental fixed effects including year, season, age at sampling, time since last treatment and last anthelmintic drug received. This was implemented for the only ponies recorded thrice a year, over at least 2 years. Using these estimates, the ten most susceptible and ten most resistant ponies were retained for this study. Median FEC over the past 5 years were 800 and 0 eggs/g for S and R ponies respectively^25^. These two groups of ponies were monitored throughout a grazing season during summer 2015.

To remove any intercurrent nematode infection, ponies were administered moxidectin and praziquantel (Equest Pramox^®^, Zoetis, Paris, France, 400 μg/kg of body weight of moxidectin and 2,5 mg/kg of praziquantel) three months before the start of the experiment and kept in-door. They were subsequently placed on a 7.44 ha pasture from mid-June to the end of October 2015. To eliminate residual egg excretion observed in some ponies, they were treated with pyrantel (Strongid^®^ paste, Zoetis, Paris, France; single oral dose of 1.36 mg pyrantel base per kg of body weight) 30 days after the onset of pasture season.

To identify blood biomarkers of infection, plasma samples were taken at 0, 24 and 132 days after the start of the trial to compare metabolomes of parasite-free horses (maintained in-door or grazing) with that of infected grazing individuals. These were matched with FEC, average daily weight gain (ADG), haematological and blood biochemistry parameters and faecal microbiota as previously outlined^25^. Nutritional information has been outlined previously^25^.

All experiments were conducted in accordance with EU guidelines and French regulations (Directive 2010/63/EU, 2010; Rural Code, 2018; Decree No. 2013-118, 2013). All experimental procedures were evaluated and approved by the Ministry of Higher Education and Research (APAFIS# 2015021210238289_v4, Notification-1). Procedures involving horses were evaluated by the ethics committee of the Val de Loire (CEEA VdL, committee number 19) and took place at the INRAE, Experimental unit of Animal Physiology of the Orfrasière (UE-1297 PAO, INRAE, Centre de Recherche Val de Loire, Nouzilly, France)

### Clinical parameters measurement and modelling

FEC were performed on 5 g of fresh faecal material after intra-rectal sampling and diluted in 70 mL of NaCl solution (d = 1.2) following a modified McMaster technique with a sensitivity of 50 eggs/g^60^.

Haematological and serum biochemical parameters were recorded at PFIE (UE-1277 PFIE, INRAE, https://doi.org/10.15454/1.5535888072272498e12). Haematological parameters were determined after 15-min stirring at room temperature with a MS9-5 Haematology Counter^®^ (digital automatic haematology analyzer, Melet Schloesing Laboratories, France). Recorded parameters encompassed erythrocyte, micro- and macro-erythrocyte counts, associated mean corpuscular volume, haematocrit, and mean corpuscular haemoglobin concentration. In addition, circulating thrombocytes were counted as well as leukocytes including lymphocytes, monocytes, neutrophils, basophils and eosinophils. For further analysis, an albumin to globulin ratio (AGR) was also considered.

In addition, serum biochemical parameters were quantified using Select-6V rings with the M-Scan II Biochemical analyser (Melet Schloesing Laboratories, France). These parameters included albumin, cholesterol, globulin, glucose, alkaline phosphatase (ALP), total proteins (TP) and urea concentrations.

Normality of haematological and biochemistry parameters was tested with the Shapiro-Wilk’s test (*shapiro.test()* function in R) and variables showing test value below 0.90, *i.e.* FEC, glucose, ALP and AGR concentrations, and eosinophil count were log-transformed (supplementary Table 4). Validation of monocyte and neutrophil counts were obtained from a group of 25 ponies in October 2020 using the same setting.

### Proton Nuclear Magnetic Resonance (^1^H NMR) data acquisition and processing

Samples were collected in heparin coated tubes. Whole blood drawn for plasma generation was refrigerated immediately at 4°C to minimize the metabolic activity of cells and enzymes and kept the metabolite pattern almost stable. After clotting at 4°C, the plasma was separated from the blood cells and subsequently cryopreserved at -80°C and shipped as a single batch for ^1^H-NMR profiling. Plasma samples were thawed on ice, 200 µl were mixed with 500 µl of deuterium oxide (D_2_O) containing sodium trimethylsilylpropionate (TSP, 1 mM). Samples were vortexed, centrifuged (5,500 g; 10 min; 4°C) and 600 µL of supernatant were transferred into 5 mm NMR tubes.

The ^1^H-NMR analysis was performed on a Bruker Avance III HD spectrometer (Bruker, Karlsruhe, Germany) operating at 600.13 MHz, and equipped with a 5 mm reversed ^1^H-^13^C-^15^N-^31^P cryoprobe connected to a cryoplatform. ^1^H NMR spectra were acquired using a Carr-Purcell-Meiboom-Gill (CPMG) spin echo pulse sequence with a 2-second relaxation delay. The spectral width was set to 20 ppm and 128 scans were collected with 32k points. Free induction decays were multiplied by an exponential window function (LB=0.3 Hz) before Fourier Transform.

Spectra were manually phase and baseline corrected using Topspin 3.2 software (Bruker, Karlsruhe, Germany). All spectra were referenced to TSP (d 0 ppm). The spectral data were imported in the Amix software (version 3.9, Bruker, Rheinstetten, Germany) to perform data reduction in the region between 9.0 and 0.5 ppm with a bucket width of 0.01 ppm. The region between 5.1 and 4.5 ppm corresponding to water signal was excluded and data were normalized to the total intensity of the spectra.

^1^H-NMR data were filtered from noise-related signals and finally consisted in 791 buckets ranging from 8.585 to 0.505 ppm (Supplementary Table 1). Buckets were annotated based on similarity of chemical shifts and coupling constants between plasma samples and reference compounds. The comparison was performed between one dimensional analytical data and reference compounds acquired under the same analytical conditions in our internal database, as well as from public databases like the Human Metabolome Database (HMDB, http://hmdb.ca/) and the Biological Magnetic Resonance Data Bank (http://www.bmrb.wisc.edu/). Manually curation was performed to isolate buckets corresponding to the same metabolite. Intensities of these metabolite-specific buckets were summed together to define metabolite-associated signals (hereafter referred to as “signals”), yielding 119 signals for further analysis.

To prevent spurious signals linked to age differences between individuals, metabolite levels at D0 were regressed upon pony age. None of the 119 metabolite signal intensities showed variation associated with pony ages at the considered cut-off (FDR < 5%).

### 16S faecal microbiota data for data integration

Total microorganism’s DNA was extracted from aliquots of frozen fecal samples (200 mg), using E.Z.N.A.^®^ Stool DNA Kit (Omega Bio-Tek, Norcross, Georgia, USA). The V3-V4 16S rRNA gene amplification and sequencing have been described elsewhere^25^.

For data integration purpose (described in next paragraph), 16S rRNA gene sequencing data generated from these ponies were re-analysed using qiime2 v.2020.2^61^. Adapter and primers were removed from sequencing data using cutadapt v2.1^62^. Trimmed data was subsequently imported into qiime2, and denoised using the Divisive Amplicon Denoising Algorithm from dada2^63^. In this workflow, Operational Taxonomic Units (OTU, sequence cluster defined by their dissimilarity level) are replaced by so-called amplicon sequence variant (ASV) that matches observed genetic variation in bacterial 16S rRNA gene amplicons instead of relying on a clustering operation^64^. Reads were quality filtered (maximal expected error of 1 for both reads), chimera trimmed (following the default “consensus” option) and the reads trimmed (6 and 20 bp for forward and reverse reads respectively) to yield 3,054,206 reads. ASVs were subsequently filtered to retain that found in at least two individuals and supported by five reads, before alignment and phylogeny building with mafft and fasttree respectively. ASVs were subsequently assigned taxonomy using a naive Bayes classifier trained on the green genes reference database (gg_13_8) clustered at 99% similarity. This workflow left 6,208 ASVs that were aggregated to 91 genera with phyloseq (v.1.32) for subsequent analysis. Total sum scaling normalization was applied to each taxa abundances for subsequent data integration analysis.

The raw sequences of the gut metagenome 16S rRNA targeted locus are available in NCBI under the Sequence Read Archive (SRA), with the BioProject number PRJNA413884 and SRA accession numbers from SAMN07773451 to SAMN07773550.

### Statistical analysis and data integration

Our experiment focused on R and S ponies monitored during a pasture season. Using this experimental design, we interrogated their respective metabolomes 1) to characterize metabolite trajectories throughout a pasture season, 2) to identify biomarkers that would predict the pony intrinsic resistance potential, 3) to identify biomarkers that could be used to differentiate between ponies reaching FEC cut-off for anthelmintic treatment at the end of pasture season. We restricted this analysis to ponies whose predicted resistance status matched the observed FEC value at the end of pasture season, leaving nine true R (**TR**) and eight true S (**TS**) ponies respectively. Statistical analyses applied for each of these three objectives are presented below.

### Multivariate analysis of longitudinal metabolomic data in TR and TS ponies

First, we aimed to characterize metabolomic modifications occurring in TR and TS ponies throughout a grazing season. To analyse our longitudinal data, we implemented an ANOVA-Simultaneous Component Analysis (ASCA)^65,66^ with the MetaboAnalyst R package v.3.0.3^67^. This analysis first partitions the variance contained in the metabolite data across the factors of interest (susceptibility group and time point including 0, 24 or 132 days after onset of the pasture season) and their interactions, thereby correcting the data for these effects. Simultaneously, a PCA is applied to each partition for dimensionality reduction ultimately isolating the metabolites contributing most to each effect. Significance of each effect was tested by 1,000 permutations. Following ASCA, 37 outlier metabolite signals were identified and subsequently removed from further data integration analysis leaving 319 signals for subsequent analysis.

### Differential analysis of signal metabolite intensities between TR and TS ponies and between ponies in need of treatment

Second, we aimed to identify biomarkers that would either be predictive of the intrinsic resistance potential of an individual (TR vs. TS comparison at day 0) or would distinguish between individuals in need of treatment (TR vs. TS comparison at day 132). To fulfill these two aims, we respectively performed Student’s *t* tests on the 319 retained metabolite signals between TR and TS pony baselines or between individuals showing FEC below or above 200 eggs/g at day 132. To account for multiple testing, nominal *P*-values were adjusted using the Benjamini-Hochberg correction (FDR) as implemented in the *p.adjust* function (stats package v.4.0.2). Spearman’s correlations were estimated using the *rcorr* function from the Hmisc package v.4.4-1^68^.

### Data integration to identify biomarkers predictive of parasite resistance or need of treatment

As a complementary approach for biomarker identification, clinical (including blood cell population profile, blood biochemistry and average daily weight gain), metabolomic (319 signals) and faecal microbiota data were integrated to identify correlated signals associated with intrinsic resistance potential or need of treatment. This approach also has the potential to uncover biological signals that would be missed when considering each dataset independently^33^. We ran two analyses to extract the features from each dataset with best discriminant ability between TR and TS ponies either before the pasture season on one hand, or between TR and TS ponies in need of treatment at day 132 on the other hand.

In this analysis, ASV counts estimated from faecal microbiota data were aggregated at the genus level within each time point of interest (day 0 or day 132) using the *tax_glom* function of the phyloseq package v.1.32.0. They were then filtered to retain those reaching 5% prevalence (n = 45 and 41 at day 0 and day 132 respectively), and normalized with the centered log-ratio transformation of the mixOmics package^69^. ^1^H NMR data consisted in the 319 metabolite signal intensities retained following filtering and outlier identification with ASCA. Clinical data consisted of 18 parameters. At day 0, ADG was not considered as no variation occurred leaving 17 parameters. FEC was not considered as it was used to define the groups to be compared.

Data integration was performed following the DIABLO (data integration analysis for biomarker discovery using latent variable approaches for Omics studies) framework^70^ as implemented in the mixOmics package v.6.12.2. This algorithm implements a sparse generalized canonical correlation discriminant analysis (sGCC-DA)^33,70^. Briefly, DIABLO performs feature selection, thereby retaining the only bacterial genera, metabolite signal or clinical parameters with best discriminative ability between groups. Using this sparse dataset, DIABLO then seeks for latent components (linear combinations of features from each dataset, i.e. ^1^H-NMR, faecal microbiota, and clinical parameters) that simultaneously explains as much as possible of the covariance between input datasets and the status of interest, *i.e.* pony resistance potential before or under infection^33,70^. We applied this analysis to discriminate between TR and TS ponies either before the onset of pasture season (day 0) or at pasture turnout (day 132). In each case, the number of features to be retained was determined by cross-validation analysis (10 x 5-fold) with the *tune.block.splsda()* function exploring grids of 10 to 20 genera with increments of 5, 10 to 80 metabolite signals with increments of 10 and 5 to 10 clinical parameters with increment of 1. The correlation matrix between each input datasets was determined after running a partial least square analysis using the *pls()* function^69^.

Following data integration, we applied linear regression models to quantify group and time variation in the levels of seven features that distinguished between TR and TS ponies both at day 0 and day 132. These most discriminant features had either an absolute contribution of 0.25 on the first sGCC-DA axis. For metabolite signal intensities, no metabolite matched this condition and all features contributing to the 1st axis at day 0 and 132 were retained.

### Enrichment analysis

To isolate biological pathways associated with discriminant ^1^H NMR signals and bacterial taxa, enrichment analyses were run using the MetaboAnalyst v5.0^67^ and MicrobiomeAnalyst^67^ web interfaces respectively. Significant metabolite signals were tested for enrichment against KEGG annotated metabolites and disease related blood biomarkers, while enrichment analysis on discriminant bacterial genera were run using taxa collections associated with aging or disease. Any enrichment with a False Discovery Rate (FDR) below 5% was deemed significant.

### Validation in an independent set of data

We validated our biological signal on microbial and metabolomic data using the data from Peachey et al.^26^ as an independent data set because they combined both 16S rRNA gene amplicon sequencing with metabolomic analyses. While they performed ^1^H-NMR on faecal material, we wanted to evaluate how our results obtained from blood ^1^H-NMR could match theirs. Raw data were retrieved and processed as ours, but following Peachey et al.’ read truncation parameters^26^. This process left 3,305,166 sequences assigned to 5,233 ASVs. ASV counts per sample and ASV taxonomic assignments from Clark et al. 2018 and Peachey et al. 2019 are available under the github repository.

Blood parameters of interest were validated in an independent set of 24 ponies with high FEC (821 eggs/g on average, ranging between 250 and 1700 eggs/g). Increased alkaline phosphatase had already been described in infected horses and was not considered for validation.

## Data availability statement

R script is available under the https://github.com/guiSalle/STROMAEQ repository and associated data matrices will be made available upon manuscript acceptance. ^1^H-NMR Data will be deposited on Metabolomicsworkbench.org upon manuscript acceptance. The raw sequences of the gut metagenome 16S rRNA targeted locus are available in NCBI under the Sequence Read Archive (SRA), with the BioProject number PRJNA413884 and SRA accession numbers from SAMN07773451 to SAMN07773550.

## Acknowledgements

This work was funded by an Institut Français du Cheval et de l’Équitation grant. We would like to acknowledge Drs. Sonia Lamandé and Lucie Pellissier for critical comments and discussion on a previous version of the manuscript.

## Author contributions

GS: analyzed the data, drafted the manuscript. CC: ^1^H-NMR data acquisition and annotation. JC, CK, JM: sampling and parasitological data acquisition. FR: pony management and sampling. MR, NP: haematological and biochemistry data acquisition. AB: raised funding, project management. NM: analyzed the data, drafted the manuscript.

## Competing interest

The author(s) declare no competing interests.

## Notes

### Competing Interest Statement

The authors have declared no competing interest.

