## Supplementary Information for "Integrative biology defines novel biomarkers of resistance to strongylid infection in horses"

### Supplementary material

Supplementary Table 1. 1H-NMR metabolite peak intensity recorded for each individual pony 2

Supplementary Table 2. Enrichment analysis for KEGG pathway and disease-associated signatures based on metabolite peaks and faecal bacterial genera 2

Supplementary Table 3. Linear regression summary of core discriminant bacterial genera upon sampling day and experimental group 2

Supplementary Figure 1. Time-dependent metabolite peaks distribution in TR and TS ponies 3

Supplementary Figure 2. Differential metabolites between resistant and susceptible ponies under strongylid infection at day 0 4

Supplementary Figure 3. Heatmap of features best discriminating between resistant and susceptible ponies after natural infection by strongylid (day 132) 5

Supplementary Figure 4. First axis associated feature loadings following sparse generalized canonical correlation discriminant analysis (sGCC-DA) between resistant and susceptible ponies after natural strongylid infection (day 132) 6

Supplementary Figure 5. Circos plot showing features best discriminating between resistant and susceptible ponies before onset of the pasture season 7

Supplementary Figure 6. First axis associated feature loadings following sparse generalized canonical correlation discriminant analysis (sGCC-DA) between resistant and susceptible ponies before infection (day 0) 8

Supplementary Figure 7. Heatmap of features best discriminating between resistant and susceptible ponies before natural infection by strongylid (day 0) 9

Supplementary Figure 8. Core discriminant features consistent across infection conditions 10

**Supplementary Table 1. $^{1}$H-NMR metabolite peak intensity recorded for each individual pony**

See attached csv file.

For every annotated metabolite peak, the corresponding metabolite name, chemical shift is listed and measured peak intensity is given for every pony enrolled in the study. In that case, each column name is formatted as Pony_day_Group where day is day 0 or day 132 and Group is either resistant (TR) or susceptible (TS).

**Supplementary Table 2. Enrichment analysis for KEGG pathway and disease-associated signatures based on metabolite peaks and faecal bacterial genera**

See attached xlsx file.

Significant enrichment found based on metabolite peaks or bacterial genera are listed. To isolate biological pathways associated with discriminant $^{1}$H NMR peaks and bacterial taxa, enrichment analyses were run using the MetaboAnalyst v5.0 and MicrobiomeAnalyst web interfaces respectively. Significant metabolite peaks were tested for enrichment against KEGG annotated metabolites and disease related blood biomarkers, while enrichment analysis on discriminant bacterial genera were run using taxa collections associated with aging or disease. Any enrichment with a False Discovery Rate (FDR) below 5% was deemed significant.

**Supplementary Table 3. Linear regression summary of core discriminant bacterial genera upon sampling day and experimental group**

See attached .csv file.

For every bacterial genus, the effect of sampling day, experimental group and their interaction is given with corresponding standard error, *t*-test value, P-value and FDR.

**Supplementary Figure 1. Time-dependent metabolite peaks distribution in TR and TS ponies**

For each metabolite peak, a boxplot of its intensity is plotted across timepoints (day 0, 24, 132 after onset of the grazing season), with underlying data points colored by true pony susceptibility group (purple: true resistant, TR; green: true susceptible, TS).

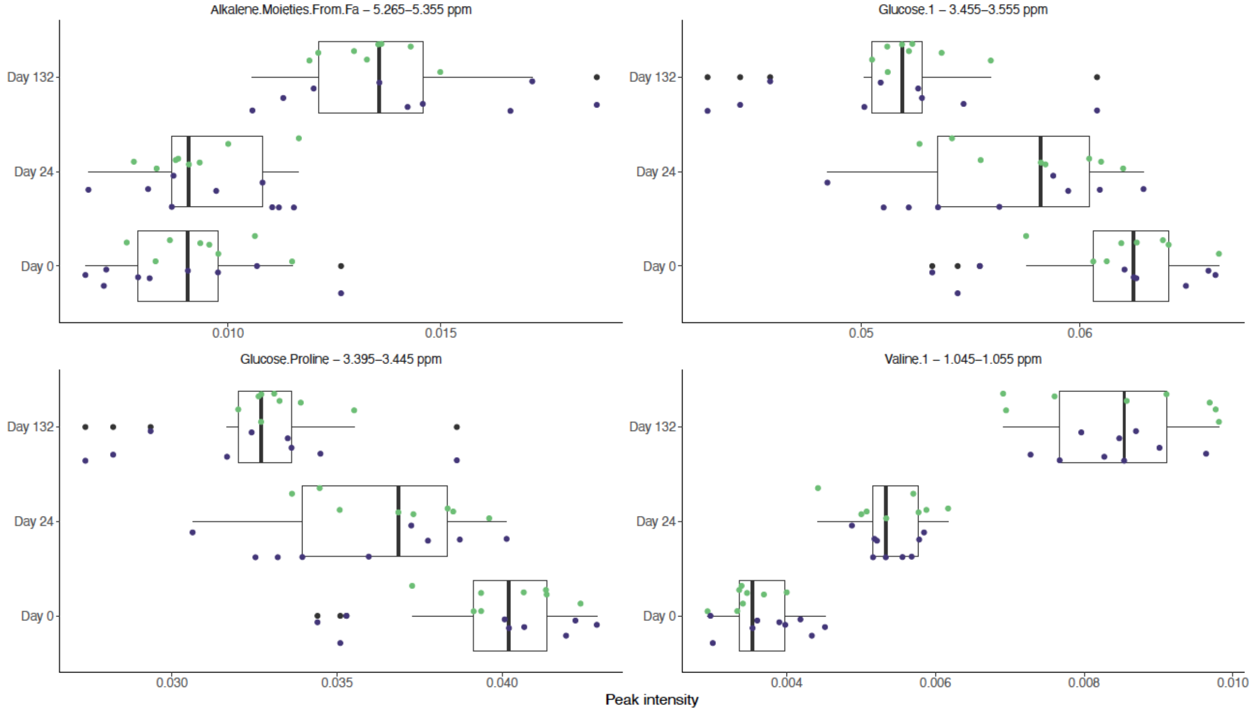

**Supplementary Figure 2. Differential metabolites between resistant and susceptible ponies under strongylid infection at day 0**

Panel a shows metabolite peak intensity distribution in each pony susceptibility group (purple: resistant, TR; green: susceptible, TS) before the onset of pasture season (day 0). Panel b shows the relationship between these metabolite peak intensities (X-axis) at day 0 and matching log-transformed Faecal Egg Count (Y-axis) at day 132.

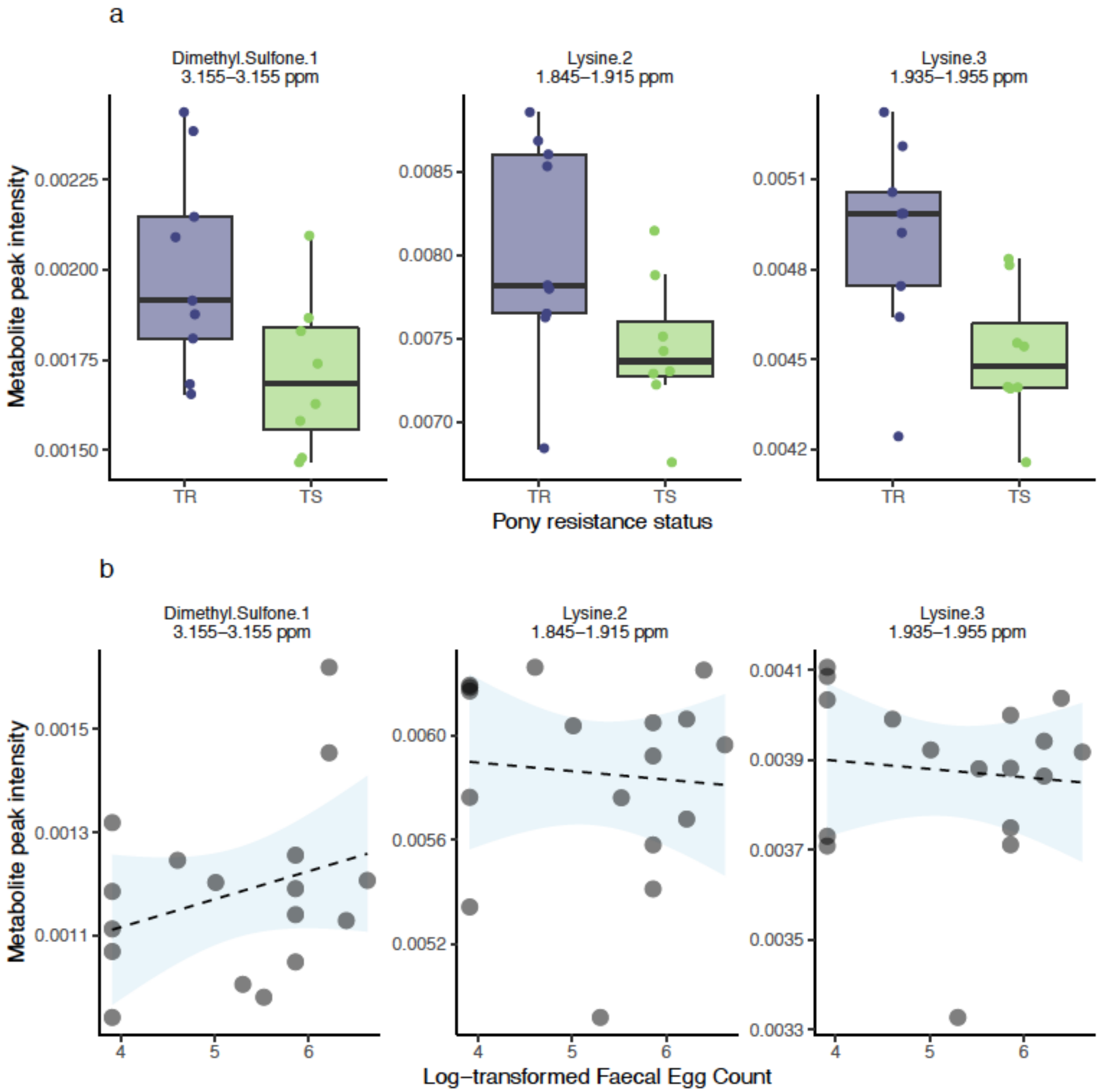

**Supplementary Figure 3. Heatmap of features best discriminating between resistant and susceptible ponies after natural infection by strongylid (day 132)**

Bacterial genera, $^{1}$H-NMR metabolite peaks and clinical parameters retained along the first axis of the sGCC-DA are clustered according to their covariation (colored from red to blue for highest and lowest values respectively). ADG: Average Daily weight Gain; ALP: Alkaline Phosphatase; MGV: Mean Globular Volume; TP: Total Proteins.

The figure highlights the opposite relationship between a core of ADG-associated genera (including *Prevotella*) and another core linked to lysine, phenylalanine and tryptophan metabolites , haematocrit and leukocyte/neutrophil counts.

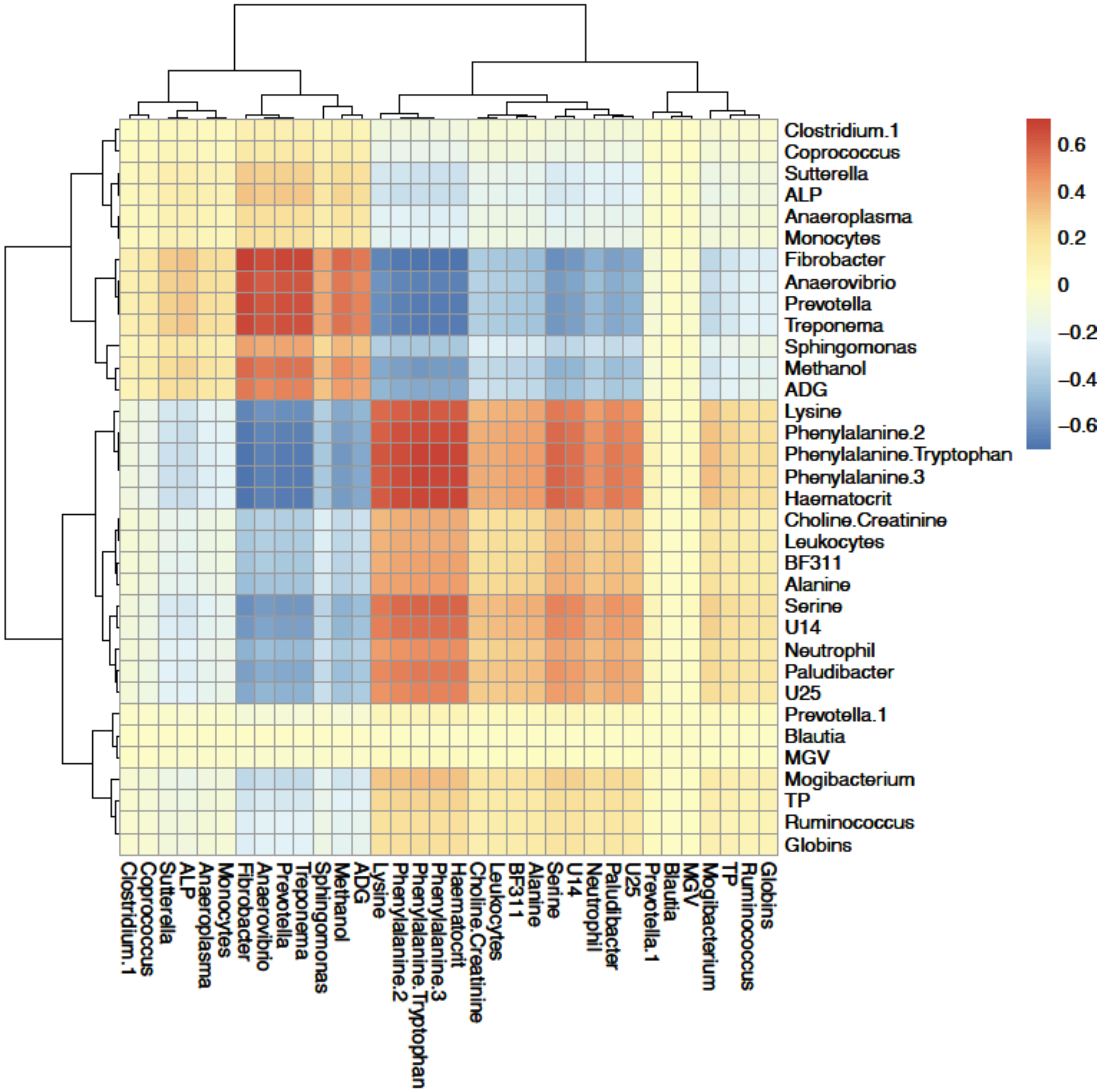

**Supplementary Figure 4. First axis associated feature loadings following sparse generalized canonical correlation discriminant analysis (sGCC-DA) between resistant and susceptible ponies after natural strongylid infection (day 132)**

For each feature group (bacterial genera, [1]H-NMR metabolite peak or clinical parameter), the respective feature loading along the first axis of the sGCDA between TR and TS ponies at day 132 (end of pasture season) is represented. Colours represent enhanced feature level in TR (purple) or TS (green) ponies respectively. ALP: Alkaline Phosphatase; MGV: Mean Globular Volume; TP: Total Proteins.

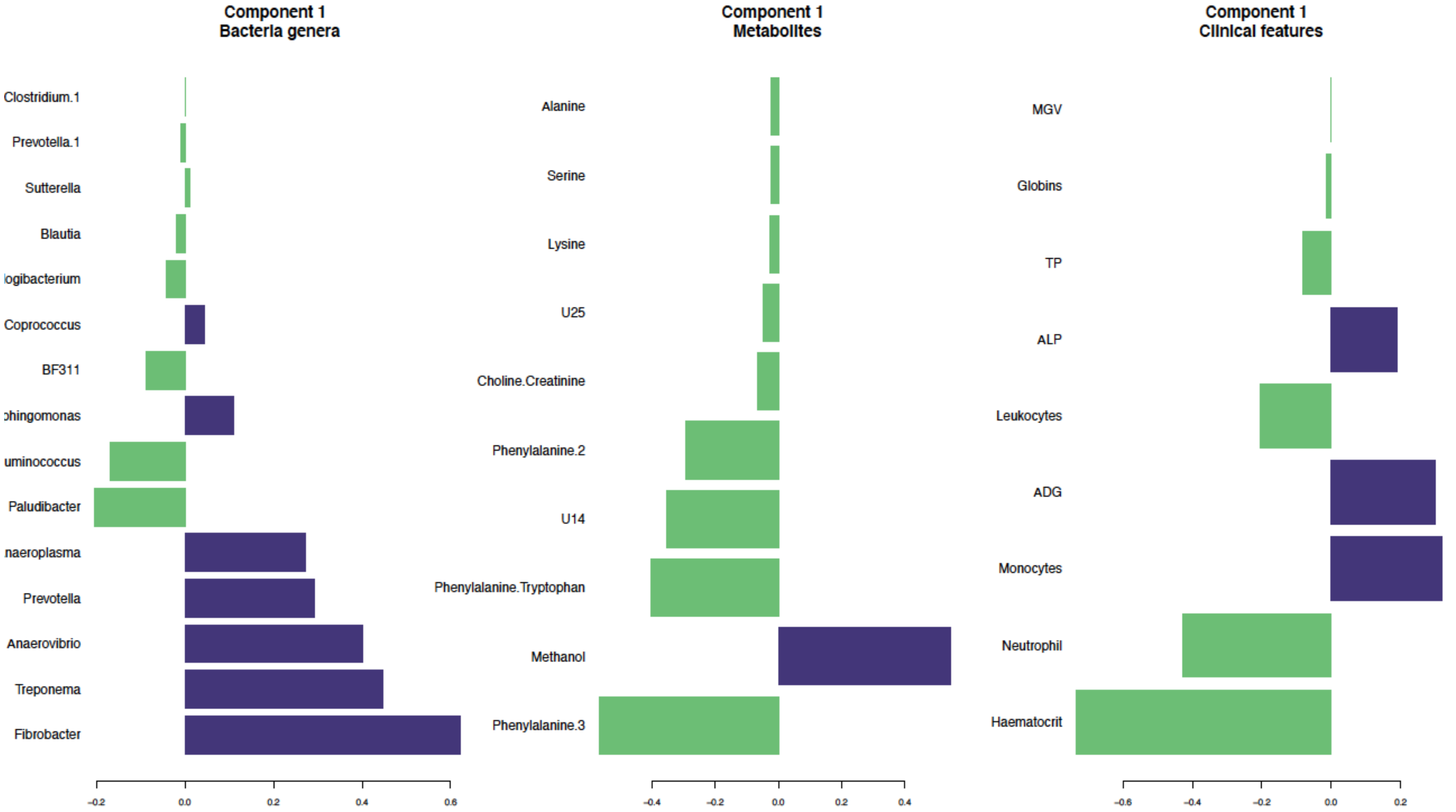

**Supplementary Figure 5. Circos plot showing features best discriminating between resistant and susceptible ponies before onset of the pasture season**

For each of the three input matrices (clinical data in yellow, bacterial genera in blue and metabolite peaks in green), features contributing to the first axis are listed, and linked if their shared correlation is above 0.5 (chocolate if positive, grey otherwise). External lines represent the feature abundance in each pony susceptibility group (resistant, TR: purple; susceptible, TS: green).

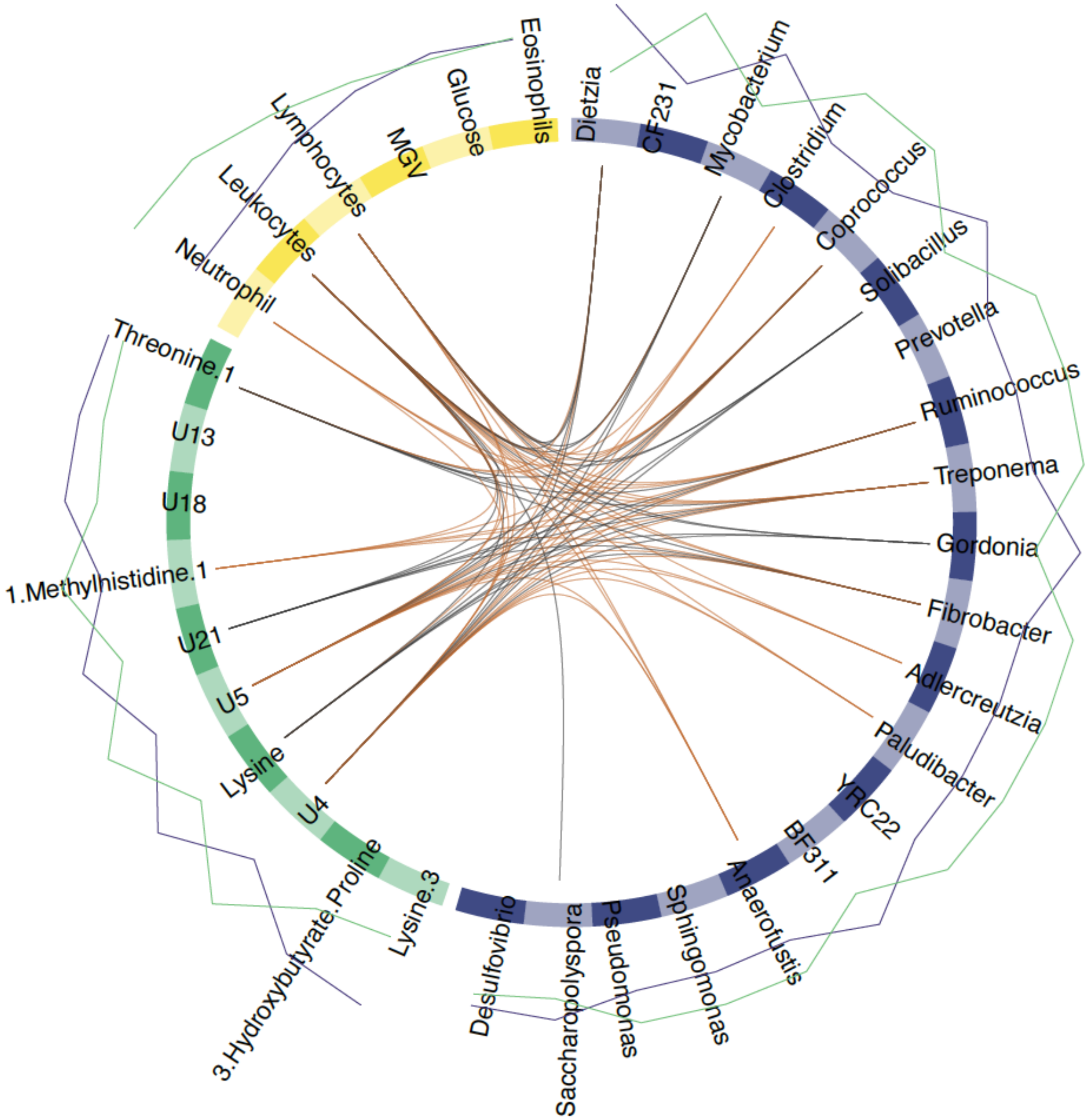

**Supplementary Figure 6. First axis associated feature loadings following sparse generalized canonical correlation discriminant analysis (sGCC-DA) between resistant and susceptible ponies before infection (day 0)**

For each feature group (bacterial genera, $^{1}$H-NMR metabolite peak or clinical parameter), the respective feature loading along the first axis of the sGCDA between TR and TS ponies at day 0 (before natural strongylid infection) is represented. Colours represent enhanced feature level in TR (purple) or TS (green) ponies respectively.

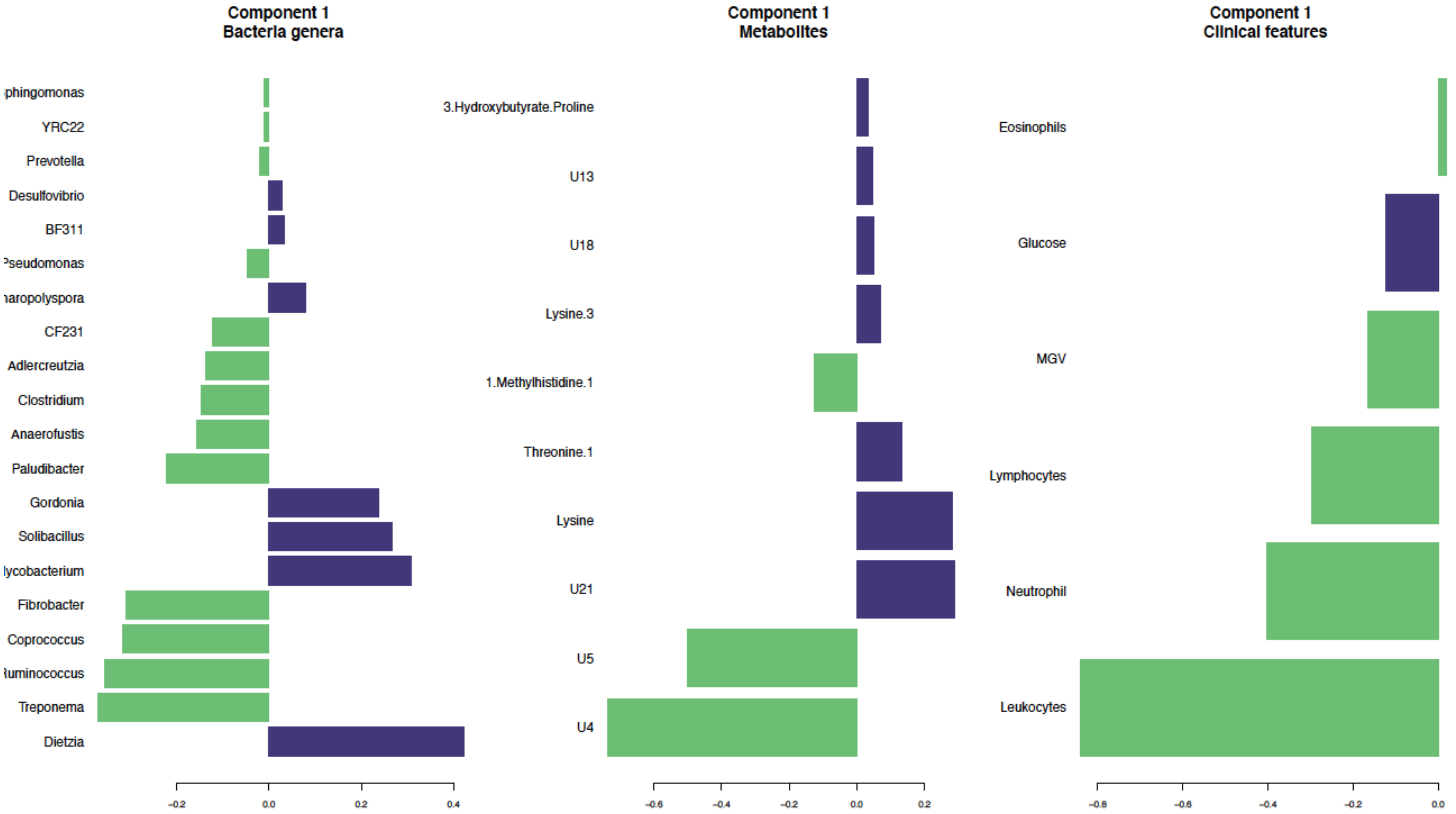

**Supplementary Figure 7. Heatmap of features best discriminating between resistant and susceptible ponies before natural infection by strongylid (day 0)**

Bacterial genera, $^{1}$H-NMR metabolite peaks and clinical parameters retained along the first axis of the sGCC-DA are clustered according to their covariation (colored from red to blue for highest and lowest values respectively). The picture highlights two blocks of strongly correlated features including a few *Corynebacteriales* species (Mycobacterium, Gordonia and Dietzia) that were negatively associated with leukocytes (lymphocyte and neutrophil) counts and butyrate-producing bacterial genera (*Anaerofustis, Coprococcus, Ruminococcus*).

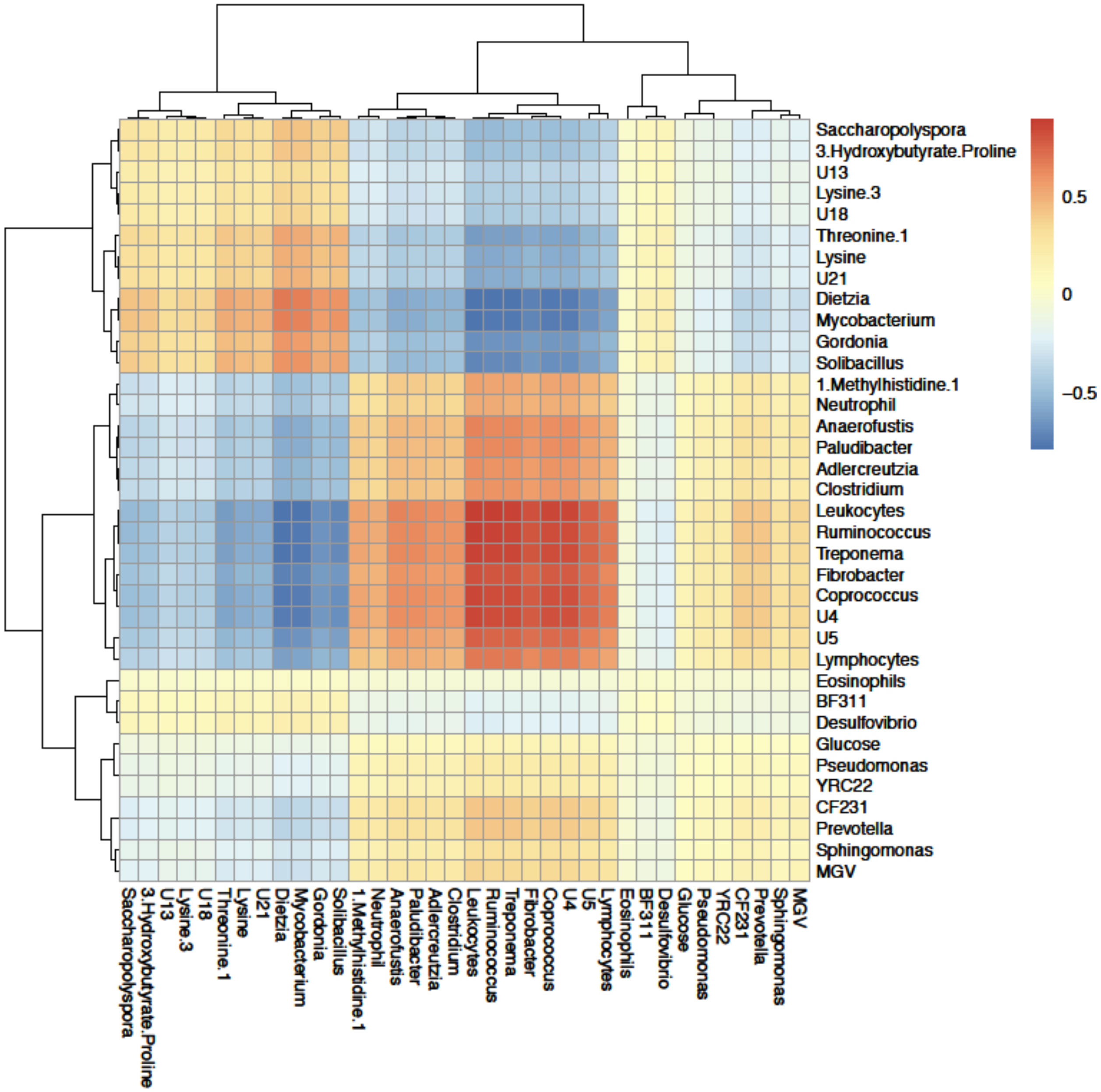

**Supplementary Figure 8. Core discriminant features consistent across infection conditions**

Measured values of seven core discriminant features between resistant and susceptible individuals are plotted across susceptibility groups (TR: resistant; TS: susceptible) and time point (day0: before infection; day132: under strongylid infection, 132 days after the onset of grazing season).

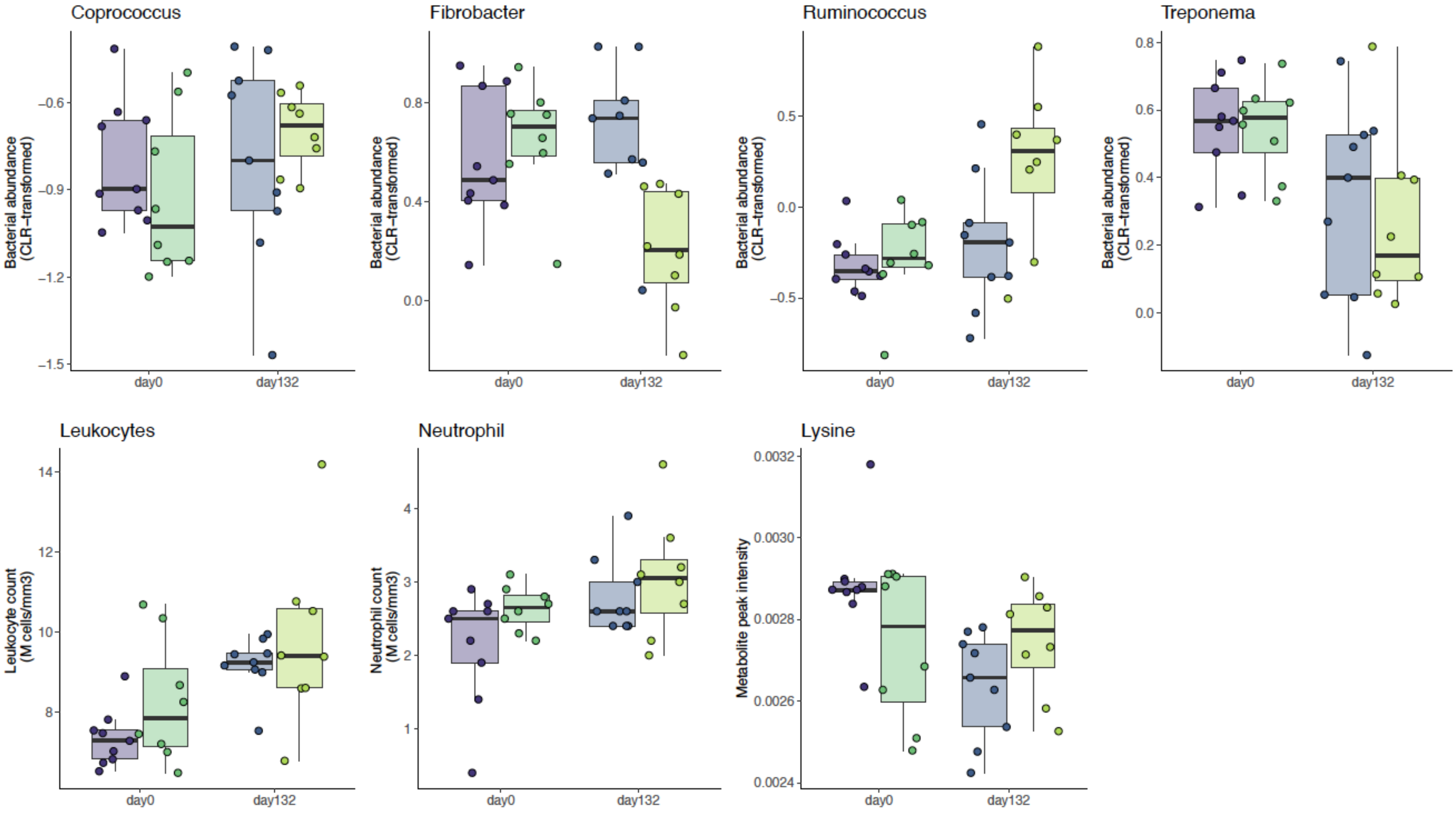
